# Forest structure and degradation drive canopy gap sizes across the Brazilian Amazon

**DOI:** 10.1101/2021.05.03.442416

**Authors:** Cristiano Rodrigues Reis, Toby. D. Jackson, Eric Bastos Gorgens, Ricardo Dalagnol, Tommaso Jucker, Matheus Henrique Nunes, Jean Pierre Ometto, Luiz E. O. C. Aragão, Luiz Carlos Estraviz Rodriguez, David. A. Coomes

**Affiliations:** Departamento de Ciências Florestais, Universidade de São Paulo, Campus “Luiz de Queiroz”. Av. Pádua Dias, nº11. Piracicaba - SP, Brazil, CEP 13418-900; Plant Sciences, University of Cambridge, Downing Street, Cambridge, UK, CB2 3EA; Departamento de Engenharia Florestal, Universidade Federal dos Vales do Jequitinhonha e Mucuri, Campus JK. Rodovia MGT 367 - Km 583, nº 5.000. Alto da Jacuba, Diamantina, Minas Gerais, Brazil. CEP 39100-000; Earth Observation and Geoinformatics Division, National Institute for Space Research (INPE); São José dos Campos, São Paulo, 12227-010, Brazil; School of Biological Sciences, University of Bristol, Bristol, BS8 1TQ, UK; Department of Geosciences and Geography, University of Helsinki, Helsinki, 00014, Finland; Impacts, Adaptation and Vulnerability Division, National Institute for Space Research (INPE). Av dos Astronautas, 1758. CEP 12.227-010. São José dos Campos, SP. Brazil; College of Life and Environmental Sciences, University of Exeter, Exeter, United Kingdom, EX4 4RJ

**Author notes:** Correspondence authors: Cristiano Rodrigues Reis. Universidade de São Paulo. Av. Pádua Dias, nº11. Departamento de Ciências Florestais, Campus “Luiz de Queiroz”. Piracicaba - SP, Brazil, CEP 13418-900., D. A. Coomes. Plant Sciences, University of Cambridge, Downing Street, Cambridge, UK, CB2 3EA.

## Abstract

Canopy gaps are openings in the forest canopy resulting from branch fall and tree mortality events. Light reaches the lower layers of the canopy through these gaps, enabling understory trees to grow and maintaining the high heterogeneity and biodiversity of tropical forests. The size-frequency distribution of canopy gaps follows a power-law distribution, and the slope of this power-law (α) is a key indicator of forest structure and dynamics.

We detected canopy gaps using a unique LiDAR data set consisting of 650 transects randomly distributed across the Brazilian Amazon Forest providing an unprecedented perspective on forest structural variation over 2500 km^2^ of forest. We then investigated how α varied with forest structure, elevation, soil fertility, water deficit, wind gust speed and lightning intensity.

We found that human modified forests had more large gaps than intact forests. Within the intact forests we observed a large-scale Northwest to Southeast pattern in α (more large gaps in the Southeast), which aligns with recent work on tree mortality rates. The two most important variables in predicting α were median canopy height and maximum modeled height, which had opposite effects on the number of large gaps. Forests with higher median canopy height contain fewer large gaps but the presence of very tall trees was associated with more large gaps. Environmental variables were of secondary importance in our model, with larger gaps occurring in drier forests with high soil fertility, wind speed, and lightning intensity.

The distribution of large gaps in the forest canopy varies substantially over the Brazilian Amazon as a result of canopy structure and mortality rates. We mapped this variation and found more large gaps in human modified forests, forests on fertile soils and those exposed to higher wind, lightning and drought stress. Increasing extreme weather events due to climate change may therefore increase the number of large gaps in currently intact forests, causing them to resemble human modified forests.

## Introduction

Gaps in tropical forest canopies arise from tree mortality and play an important role in forest regeneration processes and forest biodiversity by creating habitat heterogeneity for forest dwelling organisms (Grubb 1977, Brokaw 1985, Yamamoto 1992, Muscolo et al. 2014). Many understory plants survive in a low-light environment and depend upon these occasional gaps to capture light and grow (Marthews et al. 2008). Small gaps favour species which are shade-tolerant, while large gaps favour light-demanding pioneer species (Brokaw 1985, Yamamoto 1992). Gap colonization is driven by the nature of soil, plants and animals present at the moment of gap opening (Grubb 1977). The size of gaps is also linked to the mode of death - with broken/uprooted trees leaving larger gaps than standing dead trees (Esquivel-Muelbert et al. 2020), and associated to the type and intensity of the impact event. In this study we map the size and frequency of canopy gaps across the Brazilian Amazon and show how they are related to forest structure and environmental variables.

Remote sensing technologies make it possible to map canopy gaps over large areas of tropical forests (Lobo & Dalling 2013, Asner et al. 2013, Espírito-Santo et al. 2014, Kent et al. 2015, Wedeux & Coomes 2015, Goodbody et al. 2020, Dalagnol et al. 2021). Several studies using airborne lidar datasets have found that gap size-frequency distributions follow a simple power-law function (*f*(*x*) = *cx*^− *α*^) in which small gaps heavily outnumber large gaps in all forest environments (Kellner & Asner 2009, Asner et al. 2013, Lobo & Dalling 2013, Espírito-Santo 2014, Silva et al. 2019). Identifying power-law distributions for ecological features such as canopy gaps provides insight into the nature of gap formation processes such as tree mortality (Goodbody et al. 2020). The power-law scaling coefficient α has been associated with the type and degree of disturbance in forested areas at the landscape and regional scales (Yamamoto 1992), and can vary from less intense disturbance events (more small gaps) to mortality of large trees or damage at the stand level (more large gaps) (Asner et al. 2013, Silva et al. 2019). Extremely large gaps are very rare and they are mainly caused by wind storms (Espírito-Santo et al. 2014, Negron-Juarez et al. 2018), fire and logging events (Broadbent et al. 2008). Conversely, canopy openings due to low tree mortality and branch falls result in small gaps and are far more common (Asner et al. 2013, Espírito-Santo et al. 2014), and account for an estimated 1.28 Pg of gross carbon losses per year over the entire Amazon region - a proportion of 98.6 % of the total carbon losses due to gap formation (Espírito-Santo et al. 2014).

The size and frequency of canopy gaps is also related to the history of anthropogenic disturbance (Kent et al. 2015). Forest recovery after a disturbance event depends on the severity of disturbance, the time since it occurred, and local environmental factors (Kent et al. 2015, Cole et al. 2014), as well as anthropogenic actions such as deforestation, logging and fires (Aragão et al. 2014). In different ecological contexts such as in an area at Gola rainforest park in Africa, Kent et al. 2015 found a higher gap fraction in logged blocks (3 - 6.3%) than in old-growth forest blocks (1 - 2.3%). Wedeux and Coomes (2015), studying a peat swamp forest in Indonesia showed that, even eight years after becoming protected for conservation, logged plots had a higher gap fraction and a higher proportion of large gaps (lower α) in comparison with an old-growth forest.

We expect gap fraction and the scaling coefficient α to vary along environmental gradients, since forest dynamics is controlled by environmental variables (Phillips et al. 2004, Quesada Phillips, Schwarz, et al. 2012). Dalagnol et al. (2021) found that gap fraction across the Brazilian Amazon was positively correlated with soil nutrients (*r* = 0.46), water deficit (*r* = 0.42), dry season length (*r* = 0.41), wind speed (*r* = 0.21), and floodplains fraction (*r* = 0.27); and negatively correlated with distance to the forest edge (*r* = -0.43) and precipitation (*r* = -0.38). In particular, gap size distribution in primary forests has been linked to broad-scale patterns such as climate, topography and soils (Goulamoussène et al. 2017, Goodbody et al. 2020), as well as wind and lightning. Mortality and turnover rates in the Amazon mainly vary along an east-west gradient coinciding with a soil fertility gradient, with higher tree mortality and turnover in the rich soils of western Amazon (Phillips et al. 2004, Quesada Phillips, Schwarz, et al. 2012, Esquivel-Muelbert et al. 2020). A large proportion of Amazonian forests have also experienced water stress by intense droughts (Marengo et al. 2018), which has increased rates of tree mortality and biomass loss (Phillips et al. 2009, Phillips et al. 2010). Wind has also been linked to higher tree mortality (Rifai et al. 2016, Negron-Juarez et al. 2018), with forests in the northwest Amazon more vulnerable to windthrows and higher tree mortality than central Amazonian forests (Negron-Juarez et al. 2018). Recent work has shown that large trees are more likely to be directly struck by lightning in tropical forests with strong influences on forest structure and dynamics (Gora, Muller-Landau,). Dalagnol et al. (2021) focused on scaling-up tree mortality estimates and did not explore the gap size-frequency distribution and its relationship with environmental factors.

Local canopy height also influences the number and size-frequency distribution of canopy gaps (Wedeux & Coomes 2015). This relationship depends on the definition of a canopy gap, i.e. whether the cutoff height is defined as a relative number to local canopy height or as a fixed value (Dalagnol et al. 2021). In addition, we expect to find larger gaps in forests with taller trees, because big trees generally crush more trees along the path when it falls and produce big gaps (Grubb 1977). Therefore, interpreting environmental effects on gap properties across heterogeneous forests can be challenging. For example, a treefall event creates a much smaller gap in a forest with a substantial understory layer, as compared to the same event in sparser forest (Leitold et al. 2018, Dalagnol et al. 2021). Furthermore, the time it takes for a gap to close depends on the surrounding canopy structure (Grubb 1977, Muscolo et al. 2014) and the size of the gap (Dalagnol et al. 2019). Canopy height is related to environmental factors, and a recent study in the Brazilian Amazon showed that the presence of very large trees is explained by wind, soil, precipitation, temperature and light availability (Gorgens et al. 2020). Therefore, we expect both canopy height and environmental factors interacting to control the size and frequency of canopy gaps, however little is known about these interactions in Amazonian forests.

In this study, we use the largest tropical forest LiDAR data set to explore the relationship between gap size-frequency distribution with environmental factors and forest structure. This data set, which was collected by the “Improving Biomass Estimation Methods for the Amazon” project, provides an unprecedented perspective on forest structural variation over 2500 km^2^ of forest. We formulated four hypotheses:

H1 We expect the size and frequency of canopy gaps to be related to the history of anthropogenic disturbance i.e. that human modified forests contain more large gaps (lower α) than intact forest.
H2 We expect the size of canopy gaps to be related to maximum tree size because big trees produce larger gaps when they die (lower α in taller forests).
H3 Within intact forests, we expect gap sizes to vary along environmental gradients, i.e. in the west and southeast in accordance with water availability and soil fertility. Specifically, we expect to find fewer large gaps (lower α) in the more productive regions due to the higher turnover rates.
H4 We also expect higher wind speeds and lightning intensity to be associated with more large gaps (lower α) due to an increased rate of disturbance. Therefore, intact forests in a region with high disturbance rates may resemble human modified forests in terms of their canopy gap size distribution.

## Methods

### LiDAR data collection and processing

The “Improving Biomass Estimation Methods for the Amazon” project collected between 2016 and 2018 a considerable amount of LiDAR transects of, at least, 375 ha (12.5 x 0.3 km) each (Almeida et al. 2019, Tejada et al. 2019). The transects were allocated in forested areas by using mask layers for primary (PRODES, Inpe, 2016) and secondary forests (TerraClass, Inpe, 2014). The flights were performed at approximately 600 m height using a LiDAR Harrier 68i sensor. The survey produced a point cloud with a minimum density of 4 points.m^-2^ (Andrade et al. 2018), based on a field of view of 45° and footprint diameter between 15 and 30 cm. The data had horizontal and vertical accuracy of ±1.0 m and ±0.5, respectively (Almeida et al. 2019, Gorgens et al. 2019, Tejada et al. 2019, Gorgens et al. 2020).

We reclassified all LiDAR points into ground, vegetation and noise, excluding noise points from further analyses. The classification of the points in LiDAR data is important to provide reliable digital terrain models (DTM) and consequent height values used to estimate forest attributes, such as volume or biomass (Leitold et al. 2015; Longo et al. 2016). Points corresponding to terrain (ground points) were isolated and interpolated by the triangulation irregular network method (TIN), generating a 1-m spatial resolution DTM. In addition, we subtracted the elevation for each vegetation point by its corresponding DTM to obtain the height (Popescu & Wynne 2004). Lastly, we applied the pit-free algorithm to create the canopy height model (CHM, Khosravipour et al. 2014) using the one highest return per grid cell and triangulated them in order to obtain a 1-m spatial resolution CHM (Figure 1). The LiDAR transects were processed using LAStools software (v. 190404, Isenburg 2019).

**Figure 1.**
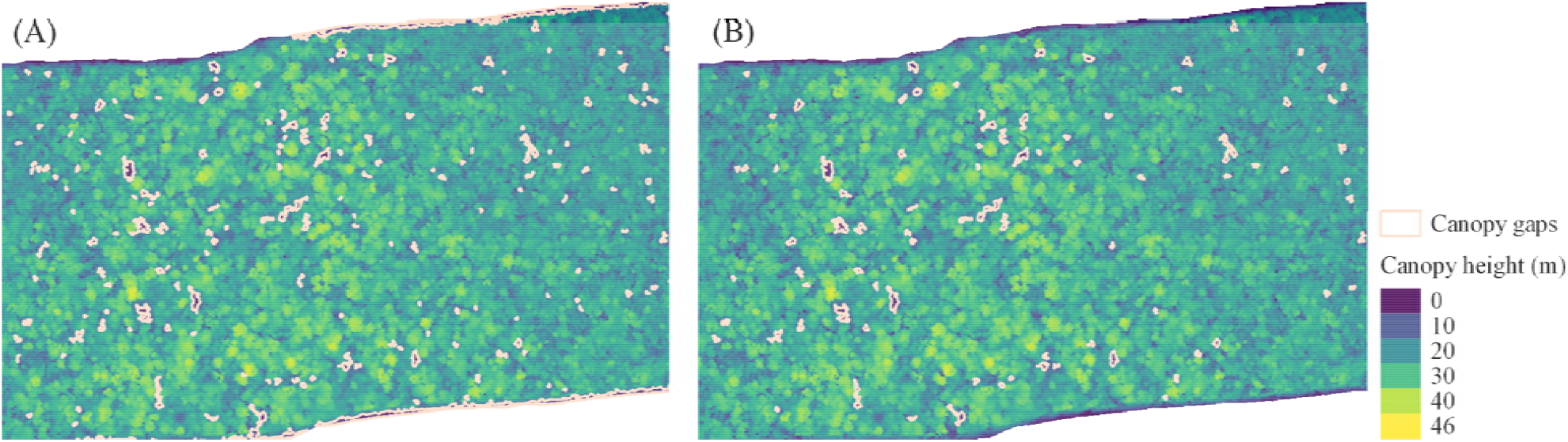
Gap filters applied in the dataset: example of gap delineation in a transect with the highest proportion of small gaps (α = 2.50) before (A) and after (B) applying the area (≥ 20 m^2^) and TPI filters.

### Extracting gaps and fitting the size-frequency distribution

To identify gaps, we took a horizontal cross-section of the CHM at 10 m above-ground and recorded agglomerations of empty pixels surrounded by full pixels (Wedeux & Coomes 2015, Silva et al. 2019, Dalagnol et al. 2021). At 10 m, Lobo and Dalling (2013) found the lowest standard error of α values compared with 2 and 5 m height. Although tradicional definition of gaps is all the polygons with areas ranging from 9 m^2^ (Wedeux & Coomes 2015) to 1 ha following previous studies, we decided to consider only gaps ranging from 20 m^2^ (Brokaw 1982, Marthews et al. 2008) to 1 ha. We also extracted the topographic position index (TPI), excluding all polygons with missing TPI values, which were associated with the transect’s edges (Figure 1 and Figure S1). We used the ForestGapR package (Silva et al. 2019) to extract all polygons within the established parameters and the spatialEco package (Evans 2020) to TPI computation.

We calculated the area of each gap to have their size-frequency and then be able to fit a simple power-law function (Equation 1):

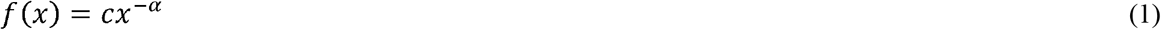

where *c* is a normalization term, x is the gap size (m^2^), and the scaling parameter α quantifies the disturbance level. As a rule of thumb, higher values of α (> 2) are found in forests dominated by small gaps and with less intense disturbance events, whereas lower α values (< 2) indicate the prevalence of more and large gaps (Asner et al. 2013, Silva et al. 2019). Deviation from the power-law pattern has also been reported at large gap sizes, but the interpretation of the scaling coefficient remains the same (Wedeux & Coomes 2015). Using the poweRlaw package (Gillespie 2015) we looked for the scaling coefficients (α) of each one of the 650 transects. We set 20 m^2^ as a minimum gap size (setXmin function) and then we estimated the power-law parameters. Finally, we plotted the α scaling coefficient to understand how α varies across the Brazilian Amazon biome.

### Delineating intact forests

To capture possible anthropogenic effects, we used the intact forest landscapes (IFL) which represent a contiguous area of natural ecosystems, showing no signs of significant human activity, and large enough that all native biodiversity could be maintained (Potapov et al. 2008). The IFL map (scale 1:1,000,000) available for 2016 was applied to divide the dataset into two categories of forests - intact and human modified forests.

We calculated the gap fraction (GF) and the number of gaps per hectare (nGAps) to perform a principal component analysis (PCA) to relate the intact and human modified forests to these three gap metrics (α, GF and nGaps). In sequence, we fitted a binomial logistic regression (Equation 2) to verify the potential of each of these metrics in distinguishing intact from human modified forest.

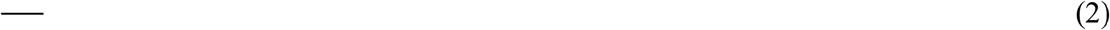

where logit(*P*) is the odds ratio, *P* is the probability of distinguishing intact forests, is the logistic regression coefficient, GF is the gap fraction in %, α is the estimated scaling coefficient from the power-law function (Equation 1), and nGaps is the number of gaps per hectare.

To test whether there are differences between the intact and the human modified forests we plotted both α distributions and ran a Wilcoxon test at 5% significance level. Due to the various possible sources of disturbances we applied further analysis only in transects within IFL. We looked for statistical differences among mean α’s by region in the Amazon (Feldpausch et al. 2011), on which we applied the post hoc Tukey’s test at 95% of confidence level.

### Environmental layers and forest structure variables

The water deficit (DEF in mm) was provided by the TerraClimate dataset, a global monthly climate and water balance for terrestrial surfaces spanning 1958–2015. With a spatial resolution of ∼5 km, this layer combined high-spatial resolution climatological normals from WorldClim with Climate Research Unit (CRU) Ts4.0 and the Japanese 55-year Reanalysis (JRA-55) data. DEF is a derived variable calculated from the difference between reference evapotranspiration and actual evapotranspiration. The reference evapotranspiration was calculated using the Penman-Monteith approach (Abatzoglou et al., 2018).

The soil cation concentration (SCC in cmol(+).kg^-1^) is a result of compiling soil data and adding indicator species to derive soil information for locations that have been sampled for plants but not soils. This approach increased the number of points to be used in soil mapping. A raster map of estimated SCC values covering all Amazonia was obtained by inverse distance weighting interpolation at the spatial resolution of 6 arcmin (∼11 km). The raster values are in log-transformed scale and represent the soil fertility gradient across the Amazon (Zuquim et al. 2019).

We used the instantaneous 10m wind gust (WG in m.s^-1^), which represents the maximum wind gust averaged over 3 second intervals, at a height of 10 metres above the surface of the Earth. This layer has a spatial resolution of ∼25 km. This variable came from the fifth major global reanalysis (ERA5) produced by the European Centre for Medium-Range Weather Forecasts (ECMWF). The reanalysis combined model data with observations from across the world into a globally complete and consistent dataset (Olauson, 2018).

The lightning frequency layer (LGT) was provided by the Lightning Imaging Sensor (LIS) with a spatial resolution of ∼11 km. The sensor collected data onboard the Tropical Rainfall Measuring Mission provided by NASA Earth Observing System Data and Information System (EOSDIS) from January 1998 to December 2013. The lightning flash rates provided the basis to detect the distribution and variability of total lightning occurring in the Earth’s tropical and subtropical regions (Albrecht et al., 2016).

The modelled maximum height (MMH in m), with a 500 m spatial resolution, was created by extracting the maximum canopy height of the same LiDAR dataset used in this study. The MMH layer was estimated by random forest approach using the distribution of giant trees across the Amazon and sixteen environmental variables as input (Gorgens et al. 2020).

The elevation was computed based on the third version of the Shuttle Radar Topography Mission (SRTM in m) provided by NASA JPL with a spatial resolution of 30 m (Farr et al. 2007; Liu et al. 2014). The SRTM mission occurred in February 2000 and collected data during ten days of operations. The digital elevation model (DEM) is available from 60° north latitude and 56° south latitude, covering 80% of Earth’s land surface. SRTM mission employed two synthetic aperture radars: a C band system (5.6 cm wavelength) and an X band system (3.1 cm).

We resampled all the layers above to a spatial resolution of 500 m using the MMH layer as reference and applying the bilinear interpolation method, cropped them to the Amazon biome extension (Figure S2), and calculated the transects median values to correlate them with their respective level of disturbance represented by α. We used the raster package (Hijmans & Etten 2012) to work with these layers. Lastly, we extracted median canopy height (MCH in m - 50^th^ percentile) to investigate how it relates to α. As an added benefit, these canopy height metrics will soon be available at a global scale from the Global Ecosystem Dynamics Inventory project (Duncanson et al. 2020) which may allow future studies to extend our predictions.

### Statistical modelling

During the exploratory analysis, we performed the Pearson correlation (*r*) among α and environmental variables and forest structure metrics. The resulting covariance matrix guided us during selection of predictor variables that should be included in the model, avoiding the strongly correlated metrics (*r* > 0.8).

We standardized all predictor variables rescaling them to have a mean of zero and a standard deviation of one. Thereafter, we fitted linear regression models to capture the variance explanation of environmental variables and forest structure metrics (Equation 3):

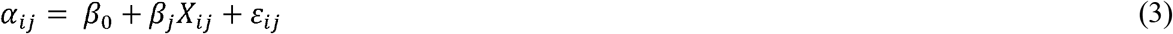

where α_i_ is the power-law scaling coefficient for transect i, β_0_ is the intercept, β_j_ is the regression coefficient for each predictor variable *X*_ij_, and ε_ij_ is the residual error.

To assess the goodness of fit we performed a graphical analysis, calculated the Akaike Information Criterion (AIC) and the root mean square error (RMSE). We also evaluated the variance inflation factor (VIF) to see how strong was the collinearity among predictors. The final model was built with the variables that most contributed to understanding α variation across the Brazilian Amazon.

## Results

### Human modified forests

We analysed 650 transects of LiDAR data evenly distributed over the Brazilian Amazon (Figure 2a) to test whether the size-frequency distribution of canopy gaps varies systematically. We split the data into intact forest and human modified forest using the intact forest landscapes product (IFL; Potapov et al. 2008). The range of α values for human modified forest (1.73 - 2.31, n = 229) overlapped with that found for intact forest (1.66 - 2.50, n = 421), although the two distributions were significantly different (Wilcoxon p-value < 0.001), which supports H1 (Figure 2b). This is illustrated by the fact that the 84% of transects (223 out of 266) with α values larger than 2 were intact forests. High values of α are associated with closed-canopy forests with a low total number of gaps and very few large gaps.

**Figure 2.**
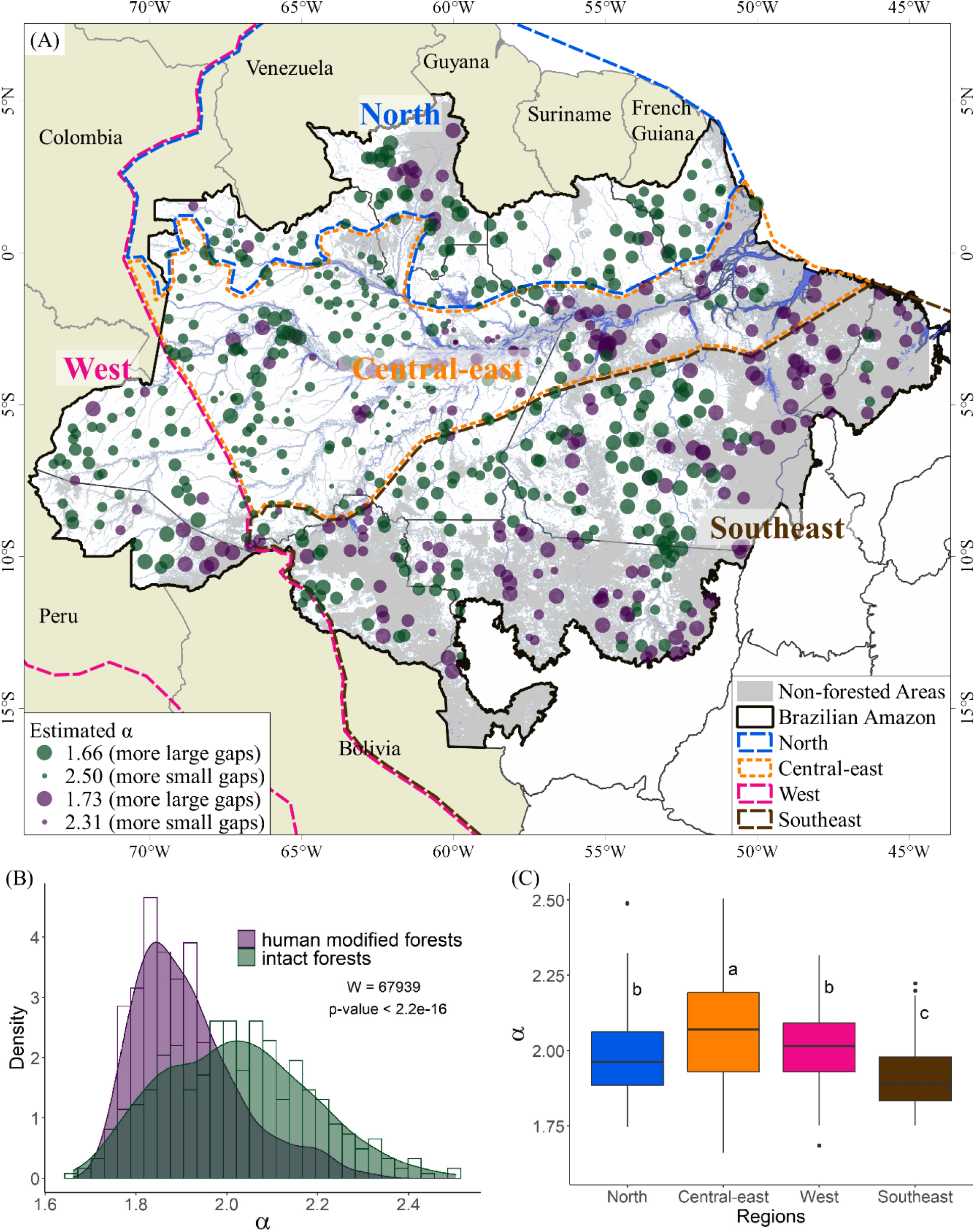
Canopy gap size-frequency distribution across the Brazilian Amazon derived from airborne lidar data (n = 650). (A) distribution of estimated power-law α’s for each transect within intact forested area (green dots, n = 421) and human modified forest (red dots, n = 229). Dashed lines represent the regions’ division obtained from Feldpausch et al. (2011); (B) different distribution of α for intact landscapes and human modified forest according to Wilcoxon test at 5% significance level; (C) Boxplot showing the α values in each region considering intact forests only. Letters show the results from post hoc Tukey’s tests comparing the mean α values among the different regions within the Amazon biome.

We found that α distinguished between intact and human modified forests whereas gap fraction and the number of gaps did not. The intact forest transects form a distinct cluster in the principal component analysis (PCA), which is characterized by high values of α (Figure 3a). Specifically, we used a logistic model to predict forest condition (intact or human modified) from the three metrics describing canopy gaps (α [β ± SE: 1.302±0.403, p-value: 0.001], gap fraction [0.686±0.762, 0.368] and the number of gaps [-0.156±0.534, 0.770]), and found that both gap fraction and the number of gaps were not significant. The gap size-frequency distribution of human modified forests had more gaps and particularly more large gaps than the intact forests (Figure 3b). Intact forests displayed a clear relationship between α and gap fraction - transects with a low gap fraction tended to have a high alpha, i.e. closed-canopy forests with few gaps and very few large gaps, whereas transects with high gap fraction had low alpha, i.e. they contain more gaps, and a greater proportion of these gaps are large (Figure S3). Human modified forests displayed a much noisier relationship between α and gap fraction (Figure S3).

**Figure 3.**
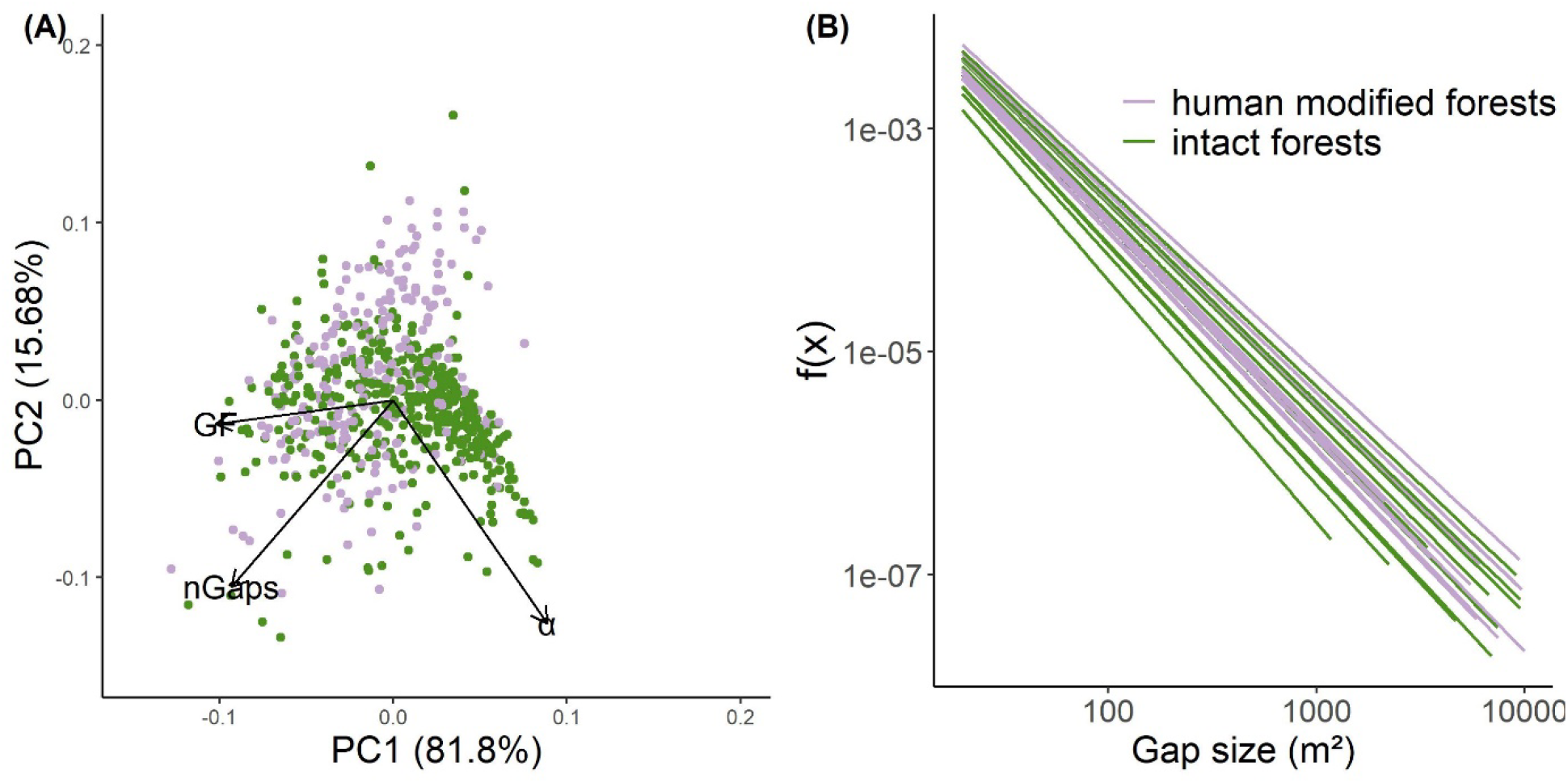
Relationship between gap metrics and forest disturbance in the Brazilian Amazon forests. (A) Principal component analysis (PCA) showing the importance of each variable in distinguishing intact and human modified forest. GF is the gap fraction (%), nGaps is the number of gaps per hectare and α is the estimated power-law scaling coefficient; (B) fitted power-law lines for a subset of transects randomly chosen for intact and human modified forests.

### Gap distributions across the Brazilian Amazon biome

These results reveal patterns of gap size-distributions across the Amazon from Northwest to Southeast (Figure 2a). Overall, we found that the Central-east region (especially Amazonas state) contained a lower number of large gaps (higher α values), whereas the South-east region (especially Pará state) showed the opposite. The South-east region had the highest proportion of large gaps (mean α ± 95% confidence interval: 1.912±0.0175). The North (1.990±0.0276) and West (1.998±0.0359) regions contained a similar distribution of gaps while the Central-east (2.058±0.0190) region had the lowest proportion of large gaps (Figure 2c).

### Drivers of variation in canopy gap size across the Brazilian Amazon

Considering only the intact forest transects, we explored the relationship between α and environmental variables as well as canopy height. Individually, we found that α was positively correlated with the median of canopy height (MCH) (*r* = 0.43), meaning that shorter forests have a higher proportion of larger gaps. This is presumably due to the chosen definition of canopy gap (an opening reaching a fixed height of 10 m above the ground) since this is more likely to occur in shorter forests. We also found that α was negatively correlated with the maximum canopy height MMH (*r* = -0.15), which suggests that the presence of large trees results in large gaps left when they die (H2).

Water deficit (DEF) was the environmental variable which displayed the strongest individual correlation with α values (*r* = -0.49, Figure 4), followed by soil cation concentration (SCC) (*r* = - 0.46). An increase in these environmental variables lead to an increase both in number and size of gaps, decreasing the estimated values of α coefficients. The natural disturbance agents, wind (WG) and lightning (LGT) were also negatively correlated with α (*r* = -0.27 and *r* = -0.13, respectively). This suggests that areas with stronger winds and more lightning have larger gaps, presumably due to disturbance (H4). The human modified forest transects’ correlation between α and environmental variables showed the same sign’s direction as the correlation for intact forests, although it was considerably weaker (Figure S4).

**Figure 4.**
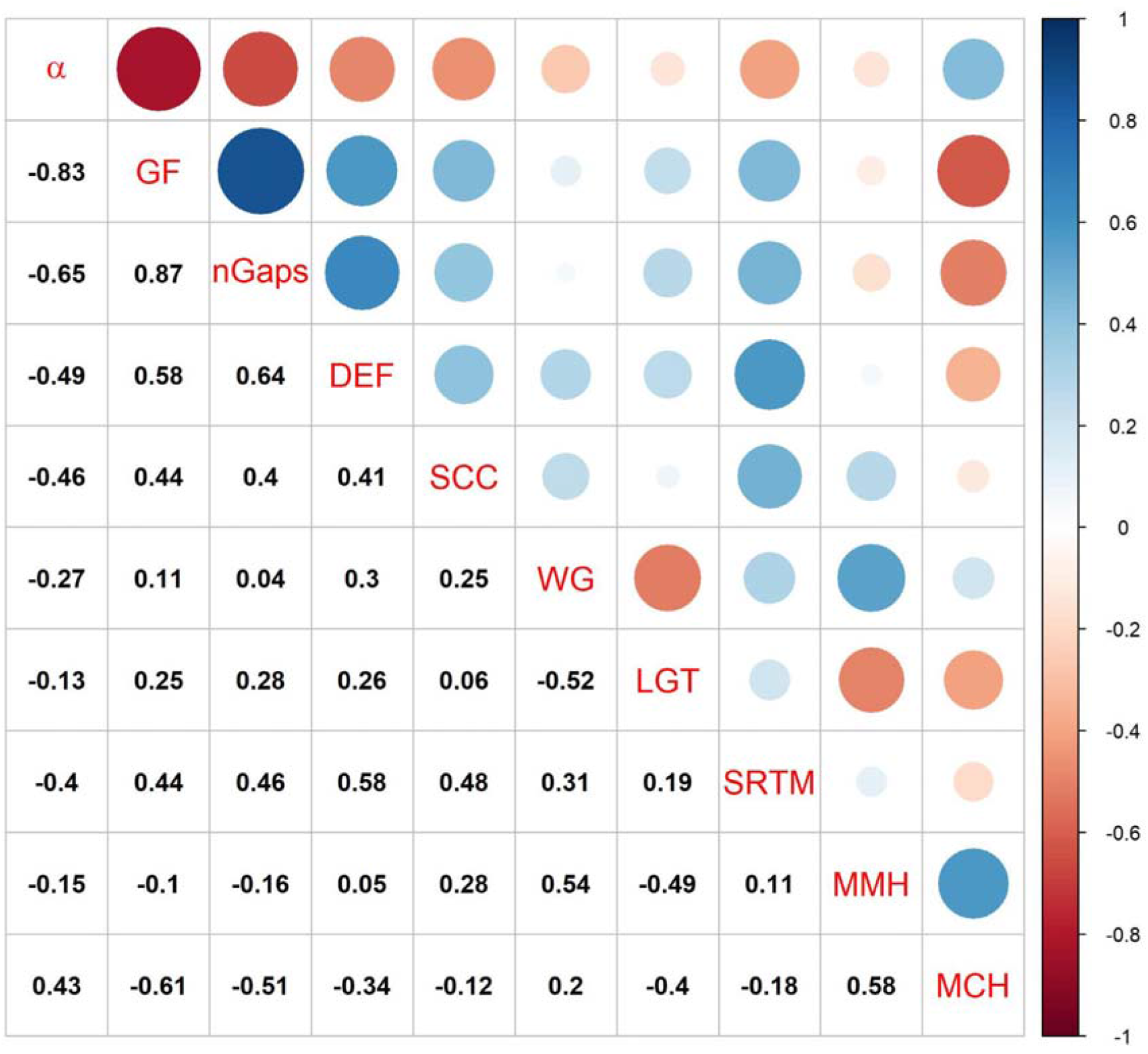
Pearson’s correlation (*r*) between LiDAR derived gap information (α, GF and nGaps) and independent variables for intact forests. α = scaling coefficient, GF = gap fraction (%), nGaps = number of gaps per hectare, DEF = water deficit (mm), SCC = soil cation concentration (cmol(+)/kg), WG = instantaneous 10m wind gust (m/s), LGT = Lightning density rates, SRTM = elevation above sea level (m), MMH = modelled maximum height (m), and MCH = median canopy height (m).

### Modelling the canopy gap size-frequency distribution

We used multiple linear regression to understand how the environmental variables and canopy height jointly explain the observed variation in gap size-frequency distribution. The best model contained median canopy height (MCH), maximum canopy height (MMH), water deficit (DEF), soil cation content (SCC), wind gust speed (WG) and lightning intensity (LGT) (Table 1). All these variables, except MCH, were associated with an increasing proportion of large gaps (i.e. a lower α). We found that, after accounting for these variables, elevation (SRTM) was not significant and not included in the model (Table S1).

**Table 1.** Power-law α coefficient fitted as function of environmental and LiDAR derived variables. Coeff = model’s β_i_ and SE Coeff = ± β_i_’s standard error. The predictor variables were: MCH = median canopy height (m) from LiDAR transects; MMH = modeled maximum height; SCC = soil cation concentration (cmol(+)/kg); DEF = water deficit (mm); LGT = Lightning density; WG = instantaneous 10m wind gust (m/s) and; VIF = variance inflation factor.

|  | Coeff | SE Coeff | t | p-value | VIF |
| --- | --- | --- | --- | --- | --- |
| | $R^2 = 0.51$; AIC = -617.02 | | | | |
| Intercept | 2.021 | 0.006 | 360.479 | < 0.001 |  |
| MCH | 0.095 | 0.008 | 11.975 | < 0.001 | 1.974 |
| MMH | -0.067 | 0.009 | -7.557 | < 0.001 | 2.494 |
| SCC | -0.029 | 0.007 | -4.340 | < 0.001 | 1.389 |
| WG | -0.026 | 0.008 | -3.187 | < 0.01 | 2.114 |
| LGT | -0.024 | 0.008 | -2.984 | < 0.01 | 2.033 |
| DEF | -0.019 | 0.007 | -2.474 | < 0.05 | 1.791 |

The two most important predictor variables were MCH and MMH, which had opposite effects on the number of large gaps. This demonstrates the importance of canopy structure as the first order effect determining gap distributions. Forests with higher median canopy height contain fewer large gaps but the presence of very tall trees is associated with more large gaps.

The environmental variables all explained similar amounts of the variation in α, after accounting for canopy height. Forests within drier areas and with more fertile soils contained more large gaps. Also, forests which experience stronger winds or more lightning contained more large gaps, presumably as a result of disturbance (H4). This model shows that much of the variation explained individually by DEF and SCC was better explained by canopy height. DEF and SCC play important roles in determining canopy height, and so their direct influence on α is substantially smaller than the pairwise correlation.

Our model performed well over most of the range of α observed in this study, but saturated at the highest values of α, which represent intact forests with very few gaps (Figure 5a). The scatter plot of residuals (Figure 5b) and the normal quantile-quantile plot (Figures 5c) showed that the residuals were non-normal distributed and heteroskedastic, as a consequence of the poor prediction of highest α values. Log transforming our data (or any other data transformation approach we test) did not improve the fit in this range. Intriguingly, these poorly predicted high alpha values were clustered in one area around Amazonas state, suggesting a difference in forest structure that our model does not account for (Figures 5d).

**Figure 5.**
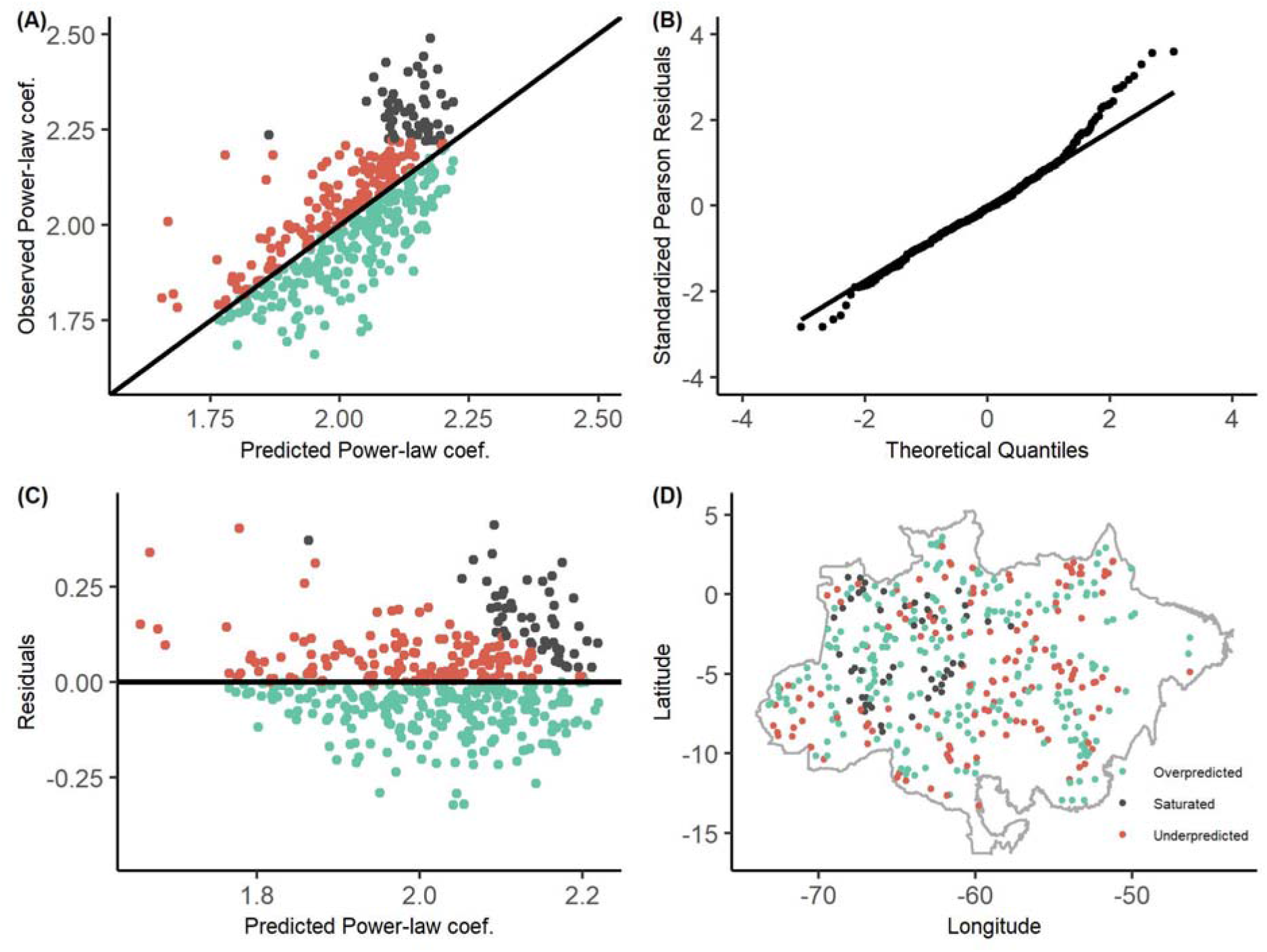
Goodness-of-fit of the final model: (A) predicted vs observed α values from the Power-law function; (B) normal quantile-quantile plot for the standardized Pearson residuals; (C) scatter plot of residuals; and (D) the distribution of the over and under predicted values for all LiDAR transects. Grey dots represent the range of poorly predicted α values (> 2.22).

## Discussion

### Human modified forests contain more large gaps

Our findings show that the α gap metric, the slope of the gap size-frequency distribution, distinguishes intact from human modified forests in the Brazilian Amazon. This is because the α highlights the disproportionate amount of large gaps in human modified forests. The remaining LiDAR gap metrics such as gap fraction and the total number of gaps were not significant in distinguishing between these two types of forests. These three metrics are commonly used to describe the observed patterns of canopy gaps and are closely related to each other. This demonstrates the importance of characterizing the size-frequency distribution when studying canopy gaps. As opposed to the number of gaps and gap fraction, α is highly sensitive to the proportion of large gaps, which is the key difference between human modified and intact forest. The high α values we found in intact forests likely indicates particularly low rates of disturbance in these areas. These high α values occurred in Central-east Brazilian Amazon within the Amazonas state, where most of the highest values of α were found (Figure 5d). All our models saturated at approximately α = 2.22 (Figure 5). This indicates that these intact forests with very few large gaps are not distinguishable by either canopy height nor the environmental variables we tested. We suggest further work to address this knowledge gap, perhaps focused on stem density, species composition or soil properties (ter Steege et al. 2006, Quesada, Lloyd, Schwarz et al. 2010, Quesada, Phillips, Schwarz, et al. 2012), for which we have particularly little data in this region.

We note that the LiDAR transects were located randomly across areas where there was forest cover - occasionally with old-growth, degraded, and secondary-growth forest mixed in the same transect (Gorgens et al. 2020). We treated each transect individually and classified it as either intact or human modified, which may have partially masked the anthropogenic effect. Dalagnol et al. (2021) also found human disturbance effects on gap fraction variability, where larger gap fractions were found over the southern and eastern Amazon regions. These regions are closer to the deforestation arch and are known for increased forest degradation. We also note that the intact forest classification measure we used also has limitations, and that a detailed knowledge of the region may allow an experienced observer to more accurately interpret the results from some of the transects.

### Large-scale trends in gap size across the amazon

We found a northwest to southeast pattern in α across the intact forests of the Brazilian Amazon. The Central-east Amazon, which is characterized by slower forest dynamics (Phillips et al. 2004, Esquivel-Muelbert et al. 2020), exhibited a higher proportion of small gaps. This contradicts our hypothesis (H3) that higher turnover leads to fewer large gaps. However, the Northern region, which also has slow dynamics (Phillips et al. 2004, Esquivel-Muelbert et al. 2020), had a lower α value (more large gaps) with a mean statistically indistinguishable from that of the Western Amazon. Phillips et al. (2004) found that the Western and Southern regions had nearly double the turnover rate (median value 2.60 %yr^-1^) compared to the Eastern and Central regions (1.35 %yr^-1^).

In addition to turnover rates, canopy gaps are also related to tree mortality rates. Tree mortality rates vary across the Amazon with a higher mortality in Western and Southern regions than in the less-dynamic Northern and Central-east regions (Esquivel-Muelbert et al. 2020). Johnson et al. (2016) found the lowest rate of stem mortality in the Central-east Amazon, followed by the Northern, Western and Southeastern regions using a network of field plots (Table 2). Dalagnol et al. (2021) predicted mortality rates using gap fraction and found a similar pattern, although with lower absolute values of mortality (Table 2). We found a higher proportion of large gaps in the Southeastern region (mean α ± 95% confidence interval: 1.912±0.0175) which aligns with previous studies and supports the relationship between gap fraction and tree mortality. However, we found a mean α for the North (1.990±0.0276) with no statistical difference from the West (1.998±0.0359) and substantially lower than the Central-east region (2.058±0.0190). This is surprising because the Northern region has previously been shown to have lower mortality rates, similar to the Central-east. We therefore conclude that these large-scale observations present a complicated picture of the relationship between canopy gaps and both turnover rates and or mortality.

**Table 2.** Mean α coefficients found in each Amazon region (Feldpausch et al. 2011) in comparison with mortality studies.

|  | Central-east | North | West | South-east |
| --- | --- | --- | --- | --- |
| Johnson et al. (2016)<br>Mortality % yr<sup>-1</sup> ± SE | $1.38 \pm 0.08$ | $1.66 \pm 0.16$ | $2.62 \pm 0.12$ | $3.19 \pm 0.38$ |
| Esquivel-Muelbert et al. (2020)<br>Mortality % yr<sup>-1</sup> (95% CI) | 1.4 (1.2–1.6) | 1.3 (1.2–1.4) | 2.2 (2.0–2.3) | 2.8 (2.4–3.4) |
| Dalagnol et al. (2021)<br>Mortality % yr<sup>-1</sup> ± SD | 0.66 ± 0.28 | 0.65 ± 0.17 | 0.8 ± 0.28 | 0.89 ± 0.2 |
| This study<br>$\alpha \pm 95\%$ CI | 2.058 ± 0.0190 | 1.990 ± 0.0276 | 1.998 ± 0.0359 | 1.912 ± 0.0175 |

One potential explanation of this disagreement between recent studies is that the regions used were designed to be allometrically distinct (Feldpausch et al. 2011), rather than being directly related to mortality. We use them to facilitate comparison with other studies (Johnson et al. 2016, Esquivel-Muelbert et al. 2020, Dalagnol et al. 2021) but our data set does not sample them evenly. In particular, the Northern and Western regions are under-represented in our sample (Figure 2a). Also, the North Brazilian Amazon contains a higher proportion of large gaps close to the savanna of Roraima, which display a different pattern of vegetation and canopy structure (Barbosa & Campos, 2007) from the rest of the Amazon.

### Accounting for forest structure

We found that α was strongly dependent on the local canopy height. We also note that canopy height is partially determined by environmental variables. Therefore, environmental variables influence α both directly (e.g. through disturbance) and indirectly through their influence on canopy height. In this study we focus on the direct effects of canopy height and environmental variables on α.

In our model of α, the two most important terms were median canopy height and maximum canopy height. Median canopy height was positively correlated with α, while maximum canopy height was negatively correlated with α. The latter effect supports our hypothesis (H2) that tall trees leave huge canopy gaps when they fall, as hypothesised by Grubb (1977). The effect of median canopy height is likely due to the fixed (10 m) cutoff height used in this study, as opposed to a cutoff height that varies with local canopy height (Dalagnol et al. 2021). Gaps in the forest canopy reaching 10 m above the ground are likely to be more common if the local canopy height is 30 m rather than 60 m. We chose a 10 m threshold since this is likely to represent mortality events, and be associated with natural regeneration, and also be relatively unaffected by ground-level noise or undergrowth (Wedeux & Coomes 2015).

### Large gaps mostly occur in productive forests

Water deficit and soil cation concentration were the individual variables most strongly correlated with α (Figure 4). However, after accounting for canopy height their importance was substantially reduced (Table 1). Our best model suggests that the direct effects of soil fertility and water deficit on α are substantially weaker than the effects of canopy height and are of a similar magnitude to the effects of wind and lightning. We note that the order of importance of the environmental variables was sensitive to the range of gap sizes we analyzed, but the orders of magnitude remained the same.

In support of H3, drier forests were associated with a higher proportion of large gaps (lower α), mainly in the Southeast fringes with frequent prolonged moisture deficits (Phillips et al. 2009). The Southern region has also a higher influence of water stress (Phillips et al. 2009). These results suggest that the primary effect of water deficit on canopy gaps is through increased mortality, especially of large trees (Nepstad et al. 2007) in drier areas, rather than through higher turnover in the more productive forests.

Soil fertility was also individually strongly correlated with α (Figure 4) and higher soil nutrient availability was associated with larger gaps. For example, Acre state has fertile soils (Quesada, Phillips, Schwarz, et al. 2012, Zuquim et al. 2019) and high productivity which results in high turnover rates (Phillips et al. 2004) and large gaps. The Southeast region also contains fertile soils (Zuquim et al. 2019), which are strongly associated with a number of important forest attributes, particularly species occurrence (ter Steege et al. 2006, Figueiredo et al. 2018, Tuomisto et al. 2019). Conversely, the poor nutrient soils found in the centre of the biome, mostly Amazonas state (Figueiredo et al. 2018), was associated with smaller gaps. Overall, our results partially support H3 i.e. that forests in drier areas had larger gaps, following similar regional patterns previously described for mortality rates (Phillips et al. 2004, Phillips et al. 2009, Quesada Phillips, Schwarz, et al. 2012, Zuquim et al. 2019).

### Wind and lightning are associated with large gaps

Increased wind gust speeds and lightning frequency were associated with more large canopy gaps (lower α) suggesting that these large gaps are caused by disturbance (H4). These effects were of a similar magnitude to those of SCC and DEF.

In extreme cases, wind disturbance can cause extensive damage (gaps >10 ha) to the forest canopy (Negron-Juarez et al. 2018, Espírito-Santo et al. 2014), but the frequency of smaller scale wind disturbance is more difficult to study. Wind may be the direct cause of death for some individual trees and will also cause damaged / dead trees to snap or uproot, increasing the size of canopy gaps (Esquivel-Muelbert et al. 2020). Individual trees acclimate to their local wind environment (Bonnesouer et al 2016) so when they are exposed to increased wind loading, for example due the creation of a nearby canopy gap, they are more likely to be damaged (Mitchell et al. 2013, Aleixo et al. 2019, Kamimura et al. 2019). This leads to a gap ‘contagion’ effect where large gaps may grow over time (Wedeux & Coomes 2015).

We also found that increased lightning frequency was associated with more large gaps. Lightning is often underestimated as a driver of tree mortality, partly because it can take many years for a tree to die (Yanoviak et al 2020) and the proximate cause of death may be mislabelled (e.g. as wind damage). Recent studies show that a single lightning strike can kill multiple trees, that it predominantly affects taller trees, and that lightning could be responsible for approximately 40% of the mortality of tall trees in lowland tropical forests (Yanoviak et al 2020; Gora, Burchfield, Muller-Landau et al., 2020).

The chronic effects of wind (Ennos 1997) and lightning (Gora, Burchfield, Muller-Landau et al., 2020) necessarily influence forest structure in the long term. For example, recent work found that low wind speeds were a key factor in determining the presence of ‘giant’ trees in the Amazon (Gorgens et al. 2020). This will have knock-on effects on the presence of large gaps as suggested in H2. It is important to consider the different time-scales of the main processes determining α. Water and nutrient gradients have long-term effects on forest structure and species composition (ter Steege et al., 2006). The immediate effects of disturbance are short-lived in the tropics since canopy gaps will close after 3-6 years due to natural regeneration (Brokaw 1985). Repeated disturbance can have long-term impacts on forest structure, but these are more difficult to predict and poorly represented by our wind layer. For instance, decades of high deforestation rates left behind a legacy of fragmentation, increased forest edges, and degraded forests (Aragão et al. 2014). It is therefore understandable that our snapshot of the size-distribution of canopy gaps, collected between 2016 and 2018, was more strongly related to the water (*r* = -0.50) and nutrient (*r* = -0.46) layers rather than the wind (*r* = -0.26) and lightning (*r* = -0.13) layers.

## Conclusions

Canopy gaps are a key aspect of forest structure and dynamics, marking the balance between disturbance and regeneration in dense tropical forests. This study provides a new understanding of the variation in canopy gaps across the Brazilian Amazon. We used a unique data set consisting of 650 LiDAR transects to characterize the gap size-frequency distribution and explore how it varies with forest structure and environmental variables.

The power-law scaling coefficient (α) was the only gap metric capable of distinguishing intact from human modified forests. As expected, human modified forests contained more large gaps than intact forests. The scaling coefficient also revealed a Northwest to Southeast gradient in gaps size, which aligns with recent mortality studies.

Within intact forests, well-structured canopies associated with high median canopy heights contained fewer large gaps, but the presence of very tall trees was associated with more large gaps. Higher soil fertility, wind speed, lightning and drier forests were associated with more large gaps. These findings show that increasing extreme weather events may cause currently intact forests to resemble human modified forests with more large canopy gaps.

## Acknowledgments

This work was supported by the Coordenação de Aperfeiçoamento de Pessoal de Nível Superior - Brasil - CAPES - Finance Code 001, Univer-sity of São Paulo (USP/ESALQ) and National Institute for Space Research (INPE). TDJ and DAC were supported by the UK Natural Environment Research Council (grant number NE/S010750/1). TDJ was supported by a UK NERC Independent Research Fellowship (grant number: NE/S01537X/1). RD was supported by the Sao Paulo Research Foundation (FAPESP, grant number 2019/21662-8). MHN was supported by the Academy of Finland (decision number 319905). L.E.O.C.A. was supported by CNPq (processes 305054/2016-3 and 442371/2019-5) and FAPESP 18/15001-6. We thank Mauro Assis for facilitating access to the LiDAR data.

## Author contributions

CRR, DAC and TDJ conceived the ideas and designed methodology; JPO provided the “Improving Biomass Estimation Methods for the Amazon’’ LiDAR data; CRR and TDJ analysed the data with support of DAC; CRR and TDJ led the writing of the manuscript, in close collaboration with DAC, EBG, RD, MHN, TJ, JPO, LCER, and LEOCA. All authors contributed critically to the drafts and gave final approval for publication.

